# Bacterial biosynthetic efficiency is constrained by cell geometry and intracellular diffusion

**DOI:** 10.64898/2026.07.25.740728

**Authors:** Arianna Cylke, Indrajit Badvaram, Shiladitya Banerjee

## Abstract

Bacterial metabolic strategies are tied to cell size and shape, yet how geometry constrains the efficiency of biomass production is not well understood. Here we develop a coarse-grained whole-cell model of bacterial physiology that couples proteome allocation, metabolic fluxes, and cell geometry to physical limits on surface area and intracellular diffusion. We define the biosynthetic energy efficiency as the fraction of ATP available from imported carbon that is invested in biomass, and find that it is non-monotonic in nutrient availability, peaking precisely at the onset of overflow metabolism. This identifies the metabolic switch as an optimal trade-off between efficient use of imported carbon and rapid growth. Perturbing cell morphology away from the empirical scaling laws shows that increasing surface area at a fixed volume raises both growth rate and efficiency, so the observed size and shape relations sit close to an efficiency optimum rather than being arbitrary. When the empirical growth laws are relaxed and geometry is treated as a free parameter, the model predicts a hard physical limit: the maximum sustainable cell size falls as the target growth rate rises. This ceiling arises from a conflict within the finite proteome budget between the cost of fast growth and the cost of large size, the latter set by the slowing of intracellular diffusion. A few physical rules thus delimit the metabolic strategies and size range available to bacterial life.

## 1. INTRODUCTION

Bacterial cells exhibit remarkable adaptability, tuning their physiology to changing environments [1–8]. This adaptation is governed by a complex interplay between gene expression, metabolic activity, and physical form [5, 9–13]. Classic “growth laws” describe empirical relationships between nutrient availability, growth rate, and cell size [1, 14–20], yet the underlying biophysical principles that shape these laws are not fully understood. A key metabolic strategy observed in rapidly growing microorganisms is overflow metabolism [21–23]. In this state, cells utilize low-yield fermentation even when sufficient oxygen is available for high-efficiency respiration. The result is that the carbon source is taken up faster than the respiratory chain can process it, forcing this excess carbon flux to “overflow” into partially oxidized byproducts, such as acetate, which are then excreted. Although excreting a usable carbon source appears energetically costly, this behavior is now understood as a rational allocation strategy: fermentation produces ATP at lower proteome cost than respiration, so cells favor it when protein synthesis capacity, rather than carbon, is limiting [24].

What remains poorly understood is how the cell’s physical architecture sets the point at which this switch becomes favorable. The cell surface area dictates the capacity for nutrient import and membrane-bound respiration, while the volume sets the total metabolic demand and the length scales for intracellular transport. The finite area of the cell envelope limits ATP synthase and electron-transport capacity, and this geometric bottleneck is a key trigger for the respiration-to-fermentation switch [25, 26]. Yet existing proteome-allocation models [24, 26–28] do not fully resolve how the physical constraints of cell size and shape feed back on metabolic strategy. Here we propose that a significant and previously underweighted cost of metabolism arises from these geometric and diffusion constraints.

This raises several questions that geometry, rather than proteome accounting alone, may answer: What sets the growth rate at which overflow becomes favorable, and are cells operating near their peak growth efficiency? What physical principles determine optimal cell size, and what are the metabolic consequences of deviating from it?

To address these questions, we develop a mechanistic flux balance framework that couples energy generation to growth regulation and cell size. Our model divides the proteome into key functional sectors— ribosomal proteins for translation, cell envelope proteins that dictate the membrane surface area, enzymes for respiration and glycolytic processes, and a residual sector for other proteome functions. We solve for the optimal allocation of these resources under steady-state conditions. By incorporating physical constraints from both cell geometry and intracellular diffusion, our model provides a mechanistic framework to explore the physical costs of metabolism.

We show that our model, calibrated with data from *E. coli*, reproduces known physiological behaviors, including proteome allocation shifts and the onset of acetate excretion. We then use it to show that energy efficiency for biosynthesis peaks precisely at the transition to overflow metabolism, framing this strategy as an optimal trade-off between nutrient efficiency and growth speed. By perturbing cell morphology away from the empirical scaling laws, we find that increasing surface area at a fixed volume improves both growth rate and efficiency, indicating that the observed size and shape relations sit close to an efficiency optimum. Finally, when the empirical growth laws are relaxed and cell geometry is optimized freely, the model predicts that the maximum sustainable cell size is inversely related to growth rate, a limit set by a conflict in the proteome budget between the cost of growth speed and the cost of size. Together, these results show how a small set of physical rules defines the metabolic strategies and size range available to bacteria.

## 2. METABOLIC FLUX BALANCE MODEL

We developed a theoretical framework to quantitatively study how physical and biochemical resource constraints shape bacterial growth and metabolism. The model is built on the principle of proteome allocation, where the cell must partition its finite protein synthesis capacity among competing functions like metabolism, biomass synthesis, and maintenance of the cell envelope. A key aspect of the model is that it connects cell structure and metabolic state: the allocation of proteome to the cell envelope determines the surface area, which in turn imposes physical constraints on nutrient import and energy fluxes.

We divide the cellular proteome into four key sectors (Fig. 1): ribosomes (mass fraction *ϕ*_*R*_), cell envelope proteins (mass fraction *ϕ*_*E*_), energy generation by glycolytic enzymes (mass fraction *ϕ*_*G*_) and the remaining proteins (mass fraction *ϕ*_0_), which are constrained by mass balance:

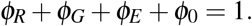

**Figure 1.**
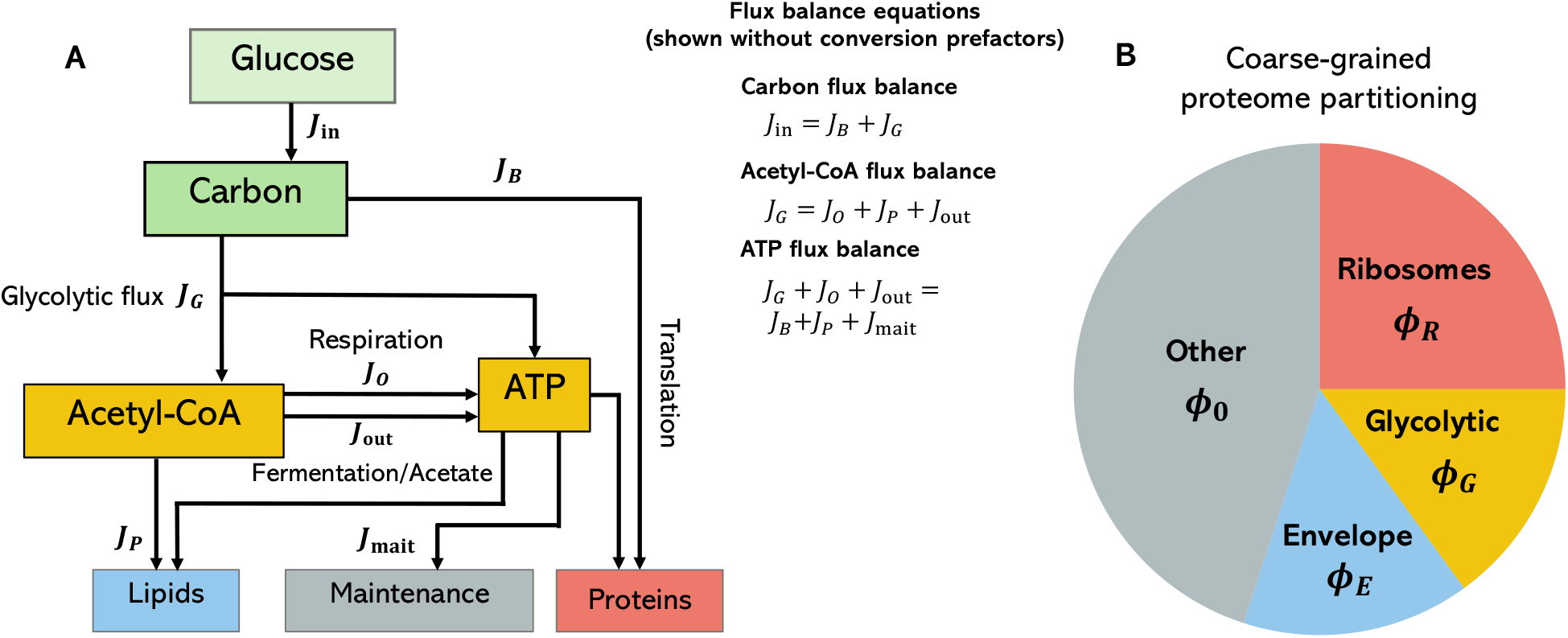
Metabolic flux balance and proteome partitioning in a growing bacterium. (A) A schematic of the flux balance model, which links proteome allocation to the geometrically-constrained metabolic fluxes that drive cellular growth. Flux balance equations are shown without conversion prefactors for brevity. See Eqs. (2)–(4) for the full flux balance equations. (B) Partioning of the cellular proteome into four coarse-grained sectors: ribosomal proteins (*ϕ*_*R*_), cell envelope proteins (*ϕ*_*E*_), energy-generating glycolytic proteins (*ϕ*_*G*_) and other proteins responsible for housekeeping, maintenance and the remaining cellular tasks (*ϕ*_0_).

The overall protein mass (*M*) dynamics, and thus the steady-state growth rate *κ*, are determined by the allocation to ribosomes:

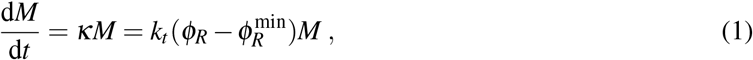

where *k*_*t*_ is the maximum translation rate and 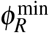 is the fraction of inactive ribosomes. The envelope proteome sector is further broken down into proteins involved in respiration *ϕ*_*O*_ (mainly ATP synthase and ETC proteins [26]) and nutrient import *ϕ*_in_, as well as membrane proteins that serve other functions *ϕ*_*E*,0_: *ϕ*_*O*_ + *ϕ*_in_ + *ϕ*_*E*,0_ = *ϕ*_*E*_. We estimate that for a typical *E. coli* cell (∼ 1 *µ*m^3^), the proteome mass fraction for nutrient import is *ϕ*_in_ = *f*_in_*ϕ*_*E*_ = 0.1*ϕ*_*E*_ (see Methods 5.1 for derivation), and use this as a fixed approximation throughout.

The cell’s metabolism is described by a set of fluxes for carbon (*C*), acetyl-CoA (*A*), and ATP (Fig. 1).

- *Nutrient Import* – Carbon enters the cell via porins on the cell surface. The import flux *J*_*in*_ is proportional to the external glucose concentration [*G*_0_] and the cell’s surface area *S* [26], which is determined by the envelope proteome fraction *ϕ*_*E*_:

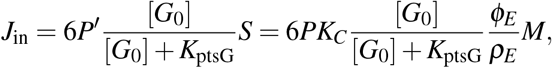

where *K*_ptsG_ = 10 *µ*M is the saturation constant of the PTS glucose transporter [29, 30], *P*^′^ = *PK*_*C*_ is the effective porin flux, *P* is the porin flux [26], *K*_*C*_ is a phenomenological flux capacity chosen as a free parameter to align with experimental overflow transition data (see Methods 5.3), and *ρ*_*E*_ is the areal density of cell envelope proteins.
- *Energy Generation* – ATP is generated through two primary pathways. Volume-based glycolytic flux (*J*_*G*_) depends on the allocation to glycolytic enzymes (*ϕ*_*G*_) and is shared between respiratory and fermentative metabolism. *J*_*G*_ spans the full conversion of glucose to pyruvate and of pyruvate to acetyl-CoA. For convenience, we refer to this combined pathway as the glycolytic pathway rather than glycolysis alone. Surface-based respiration flux (*J*_*O*_) depends on the allocation to respiratory proteins (*ϕ*_*O*_) within the cell envelope:

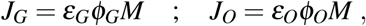

where *ε*_*G*_ and *ε*_*O*_ are the proteome efficiencies of glycolytic processes and respiration (ATP per mass per time). Crucially, respiration is geometrically constrained by the available surface area, such that *ϕ*_*O*_ ≤ *γ*(1 − *f*_in_)*ϕ*_*E*_ , where *γ* = 0.39 represents the maximum packing density [26] as derived in Methods 5.1.
- *Biomass and maintenance* – Carbon and ATP are consumed to create new biomass and for cellular maintenance. The carbon flux to biomass (*J*_*B*_) and the ATP flux for synthesizing proteins and lipids are directly proportional to the growth rate *κ*. If the carbon fraction of *E. coli* biomass is *f*_*C*_, then

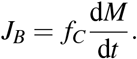

Maintenance energy flux *J*_mait_ = *aM* is a fixed cost per unit mass, where *a* gives the maintenance rate per unit of biomass [31]. We obtain the parameter *a* from the non-growth associated maintenance energy (NGAM) reported in [32].
- *Acetate Excretion* – Acetate excretion is the terminal step of the fermentative pathway. Excess acetyl-CoA can be excreted as acetate (*J*_out_), which also generates a small amount of ATP via substrate-level phosphorylation. Acetate excretion occurs only when the respiratory proteome *ϕ*_*O*_ is saturated, so *J*_out_ is zero otherwise.

The model is defined by flux balance equations for carbon (*C*), acetyl-CoA (*A*), and ATP. The change in carbon content *C* in the cell is given by the fluxes,

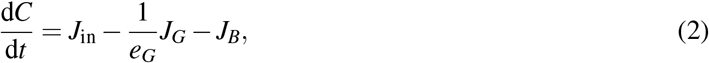

where the ATP flux from glycolytic processes (*J*_*G*_) is scaled by the number of ATP generated per carbon via the glycolytic pathway, *e*_*G*_.

Glycolytic processes convert glucose to pyruvate, which is then converted to acetyl-CoA (Fig. 1A). Acetyl-CoA is in turn used for respiration, lipid synthesis, or excretion as acetate (the conversion of each acetyl-CoA to acetate generates one additional ATP [24]):

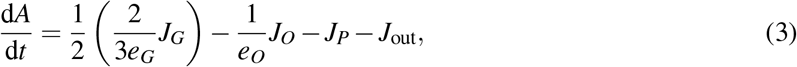

where the coefficient of 1/2 indicates 2 carbon per acetyl group, and 2/3 indicates that 1/3 carbon are expended as *CO*_2_. *e*_*O*_ denotes the ATP generated per acetyl-CoA and *J*_out_ is the rate of acetate excretion. The acetyl-CoA flux to lipid synthesis, *J*_*P*_, must be sufficient to produce new membrane for the growing cell envelope. This flux is therefore proportional to the growth rate and the envelope proteome fraction:

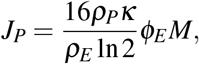

where *ρ*_*P*_ is the phospholipid surface density (see Methods 5.1).

Considering available energy from glycolysis and respiration in units of ATP, we have

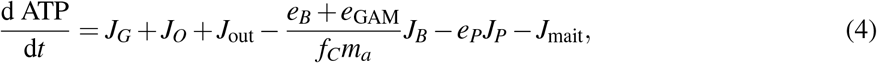

where *e*_*P*_ denotes the ATP cost per acetyl-CoA for phospholipids synthesized, *m*_*a*_ is the average mass of an amino acid, *e*_*B*_ is the ATP cost of synthesis and translation per amino acid [33], and *e*_GAM_ is the additional growth-associated maintenance energy per amino acid (estimated and normalized from [32], see Methods 5.1).

The system of equations, Eq. (1)–(4), define the dynamics of cellular growth and metabolism for a given glucose concentrations [*G*_0_] and proteome mass fractions (*ϕ*_*R*_, *ϕ*_*G*_, *ϕ*_*E*_ , *ϕ*_*O*_). To capture the cell’s distinct metabolic states, the model is implemented piecewise. For any given condition, we first test for normal metabolism by setting the acetate excretion flux to zero (*J*_out_ = 0) and solving the system for *ϕ*_*O*_. We then check if this solution is physically viable by comparing the resulting respiratory proteome (*ϕ*_*O*_) to its maximum value set by the envelope’s surface area *γ*(1 − *f*_in_)*ϕ*_*E*_ , where *γ* is a fixed ceiling on the envelope fraction available to respiratory complexes. If this constraint is satisfied, the cell remains in the normal metabolic regime. Otherwise, the cell cannot meet its energy demands through respiration alone. In this case, we implement the overflow metabolism regime by enforcing the physical limit 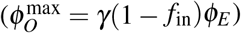 and re-solving the system for the acetate excretion flux, *J*_out_. Model parameters were estimated from experimental data on *E. coli* (Table 1), as detailed in Methods 5.1.

**Table 1.** Model parameters and variables used for our predictions unless otherwise specified for a particular figure or result.

| Parameter | Value | Physical meaning | Source |
| --- | --- | --- | --- |
| $[G_0]$ | varied | external glucose concentration | - |
| $P$ | $1.2 \cdot 10^4 \mu\text{m h}^{-1}$ | porin flux | [26] |
| $K_{\text{ptsG}}$ | $10 \mu\text{M}$ | saturation constant of PtsG glucose transporter | [29] |
| $K_C$ | $30 \mu\text{M}$ | porin associated flux capacity (phenomenological) | calibrated |
| $f_C$ | 0.47 | *E. coli* carbon fraction | [43] |
| $m_a$ | $1.1 \cdot 10^{-4} \text{MDa}$ | average amino acid mass | [44] |
| $\epsilon_G$ | $1.63\epsilon_O$ | glycolytic proteome efficiency (up to acetyl-CoA conversion) | Eq. (11) |
| $\epsilon_O$ | $1.8 \cdot 10^6 \text{ATP h}^{-1} \text{MDa}^{-1}$ | respiration proteome efficiency | Eq. (10) |
| $k_t$ | $5.9 \text{h}^{-1}$ | maximum translation rate | [2] |
| $e_G$ | 5/3 | ATP per carbon (via glycolytic pathway up to acetyl-CoA) | [24] |
| $e_O$ | 8 | ATP per acetyl-CoA (via respiration TCA) | [24] |
| $e_B$ | 23.5 | ATP cost per amino acid synthesized and translated | [33] |
| $e_P$ | 21/8 | ATP cost per acetyl-CoA (for lipid synthesis) | [44] |
| $\rho_M$ | $9.9 \cdot 10^4 \text{MDa}/\mu\text{m}^3$ | protein dry mass density | Eq. (8) |
| $\rho_P$ | $4.5 \cdot 10^6 \mu\text{m}^{-2}$ | phospholipid surface density | Eq. (12) |
| $\rho_E$ | $2.4 \cdot 10^3 \text{MDa}/\mu\text{m}^2$ | cell envelope areal protein mass density | [45] |
| $\rho_c$ | $9.9 \cdot 10^4 \text{MDa}/\mu\text{m}^3$ | crowding density scale (equal to $\rho_M$) | assumption |
| $\phi_0$ | $\geq 0.45$ | other proteome sector (housekeeping and any residual) | [2] |
| $d$ | 28.7 nm | envelope thickness | [46] |
| $\phi_R^{\text{min}}$ | 0.07 | inactive ribosome fraction | [2] |
| $a$ | $8390 \text{ATP h}^{-1} \text{MDa}^{-1}$ | non-growth associated maintenance coefficient | [32] |
| $\gamma$ | 0.39 | maximum respiration protein fraction ratio | Eq. (9) |
| $f_{\text{in}}$ | 0.1 | fraction of envelope proteome $\phi_E$ for glucose transporters | Eq. (16) |
| $e_{\text{GAM}}$ | 8 ATP/aa | growth-associated maintenance energy normalized per amino acid | Eq. (15) |

A key feature of our modeling framework is that metabolic fluxes are intrinsically linked to cell geometry. Metabolic processes such as glycolytic fluxes (*J*_*G*_) and biomass synthesis (*J*_*B*_) scale with the total protein mass *M*, which is proportional to the cell’s volume *V* via the mass density *ρ* as *M* = *ρV*. In contrast, nutrient import (*J*_in_) scales directly with the cell’s surface area *S*. While respiration (*J*_*O*_) also scales with mass, its capacity is physically limited by the surface area available for respiratory proteins, which are part of the envelope. The surface area itself is determined by the proteome fraction allocated to the envelope (*S* ∝ *ϕ*_*E*_*M*). Thus, the cell’s surface-to-mass ratio (*S/M* ∝ *ϕ*_*E*_) is a central parameter that governs the critical balance between nutrient uptake and metabolic processing.

## 3. RESULTS

### 3.1. Metabolic flux balance captures proteome tradeoffs and the switch to overflow metabolism

To investigate the model’s predictions for cellular physiology, we first constrained its geometry using the empirical growth laws for *E. coli* [15, 17], *which relate growth rate (κ*) to cell size and shape. Specifically, we used the experimentally-derived relationships between growth rate *κ* and cell volume, 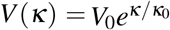 [15, 18, 19], and between volume and surface area, *S* ≈ 2*πV* ^2*/*3^ [17, 34], where *V*_0_ and *κ*_0_ are constants (see Methods 5.3). These morphological laws set the cell’s total mass (*M*) and envelope proteome fraction (*ϕ*_*E*_) for a given growth rate. Solving the steady-state flux balance equations across a range of glucose concentrations [*G*_0_] reveals that this empirically-informed model successfully captures the complex proteome allocation strategies [1, 9] and metabolic shifts observed in bacteria (Fig. 2).

**Figure 2.**
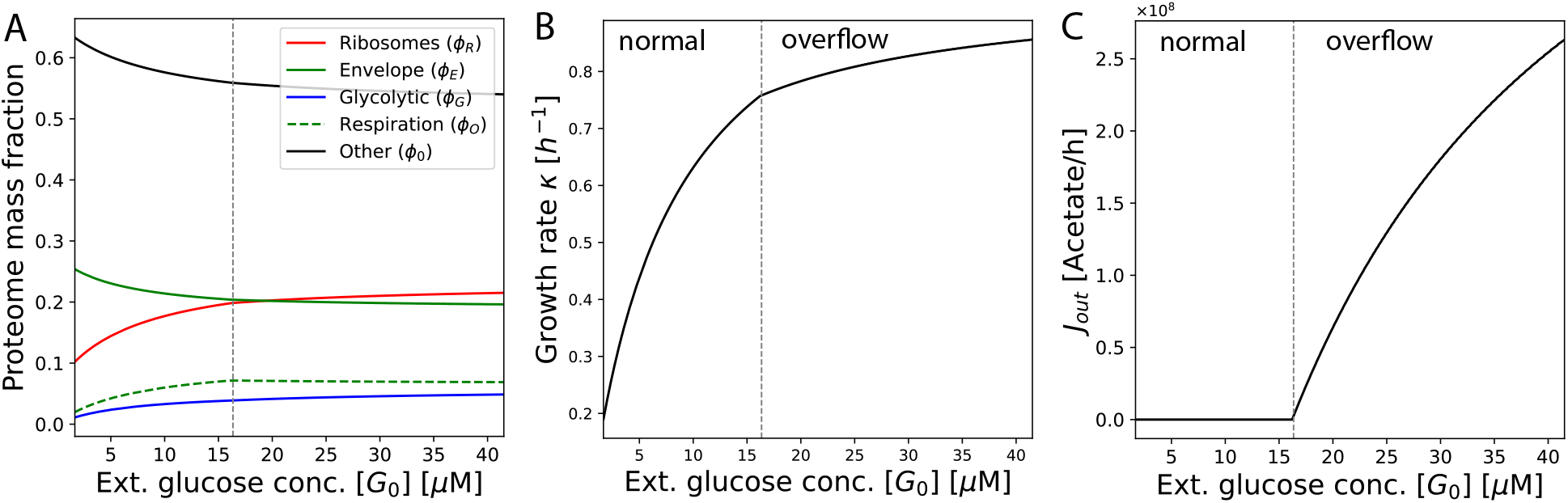
Metabolic flux balance model captures proteome trade-offs and the transition to overflow metabolism. (A) Protein mass fractions *ϕ*_*q*_ as a function of external nutrient concentration [*G*_0_], as predicted by our flux balance model. The transition point from normal metabolism to overflow metabolism is denoted with a vertical dashed line. (B) Growth rate as a function of [*G*_0_], as predicted by our flux balance model. (C) Acetate excretion as a function of [*G*_0_] as predicted by our flux balance model.

As expected, in richer nutrient environments, the proteome fraction dedicated to ribosomes (*ϕ*_*R*_) increases to support faster growth (Fig. 2A). This investment directly drives a higher cellular growth rate *κ* (Fig. 2B). Notably, the scaling of both *ϕ*_*R*_ and *κ* with nutrient availability is steeper in the initial, respiration-dominant regime compared to the subsequent overflow regime. This shift occurs because overflow metabolism is a less efficient means of converting carbon into the ATP required to fuel the ribosomal machinery.

This metabolic switch is driven by a physical bottleneck on the cell surface. While the cell initially increases its investment in the highly efficient respiratory proteome (*ϕ*_*O*_), this sector is fundamentally limited by the cell envelope fraction (*ϕ*_*E*_). As the cell grows larger in richer media, its surface-to-volume ratio decreases—a trend captured by the empirical growth laws—leading to a corresponding decrease in the overall *ϕ*_*E*_ fraction due to the trade-off with ribosomes. This results in the non-monotonic behavior of *ϕ*_*O*_, which increases initially and then marginally decreases as it hits its maximum capacity relative to the diminishing envelope mass fraction (Fig. 2A). In contrast, the glycolytic proteome (*ϕ*_*G*_), which is not surface-limited, increases steadily with [*G*_0_] to satisfy the energy demands that respiration alone can no longer meet.

The consequence of this respiratory bottleneck, as dictated by the flux balance, is the onset of overflow metabolism. We calibrated our model by comparing the onset of the overflow transition to the experimentally observed growth rate of *κ* ≈ 0.76 h^−1^ for multiple *E. coli* strains [24]. To achieve this calibration, we kept the parameters in Table I fixed, and chose a porin-associated flux capacity *K*_*C*_ such that the over-flow transition occurred close to *κ* = 0.76 h^−1^ using *e*_GAM_ = 8 ATP/aa, which we independently validated by normalizing the growth-associated maintenance (GAM) measurements reported by [32] per amino acid (see Methods 5.1 for derivation). Once overflow begins, the cell excretes acetate (*J*_out_) to balance its internal fluxes (Fig. 2C). While our model captures the qualitative behavior, it predicts a slight non-linear increase in *J*_out_ with [*G*_0_] (Fig. 2C), whereas experimental measurements appear to show a more linear relationship with the growth rate *κ* [24]. This suggests that while our core geometric and proteomic constraints are valid, secondary regulatory mechanisms may further linearize this response.

### 3.2. Optimal biosynthetic efficiency is achieved at the onset of overflow metabolism

The proteome allocation strategies predicted by our model reveal a fundamental conflict between investing in growth machinery (*ϕ*_*R*_) and maintaining cell envelope structure (*ϕ*_*E*_). This raises a critical question: what are the consequences of these trade-offs for the cell’s overall metabolic performance? To quantify this, we define the biosynthetic energy efficiency, ℰ, as the ATP invested in biomass (protein translation and lipid synthesis) divided by the total ATP that could be generated from imported carbon:

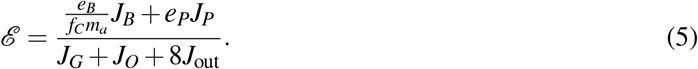

The numerator counts only the biosynthetic cost *e*_*B*_ and excludes the growth-associated maintenance over-head *e*_GAM_; the factor of 8 on *J*_out_ in the denominator reflects the opportunity cost of excreting acetate, each molecule of which represents 7 ATP foregone through respiration plus the 1 ATP recovered by substrate-level phosphorylation. Hereafter we refer to ℰ simply as the biosynthetic efficiency.

We find a non-monotonic relationship between biosynthetic efficiency ℰ and growth rate (Fig. 3A). Initially, ℰ increases as the growth rate increases. In this regime, the primary loss of biosynthetic efficiency occurs from the fixed cost of maintenance metabolism (*J*_mait_), which becomes a progressively smaller fraction of the total energy budget as the flux towards biomass, *ϕ*_*R*_, increases with increasing [*G*_0_]. While the mass fractions responsible for the ATP production terms, *ϕ*_*G*_ and *ϕ*_*O*_, are also increasing with [*G*_0_] in normal metabolism, their fluxes are balanced by Eq. (4) and only the ratio between biomass synthesis and maintenance costs is relevant.

**Figure 3.**
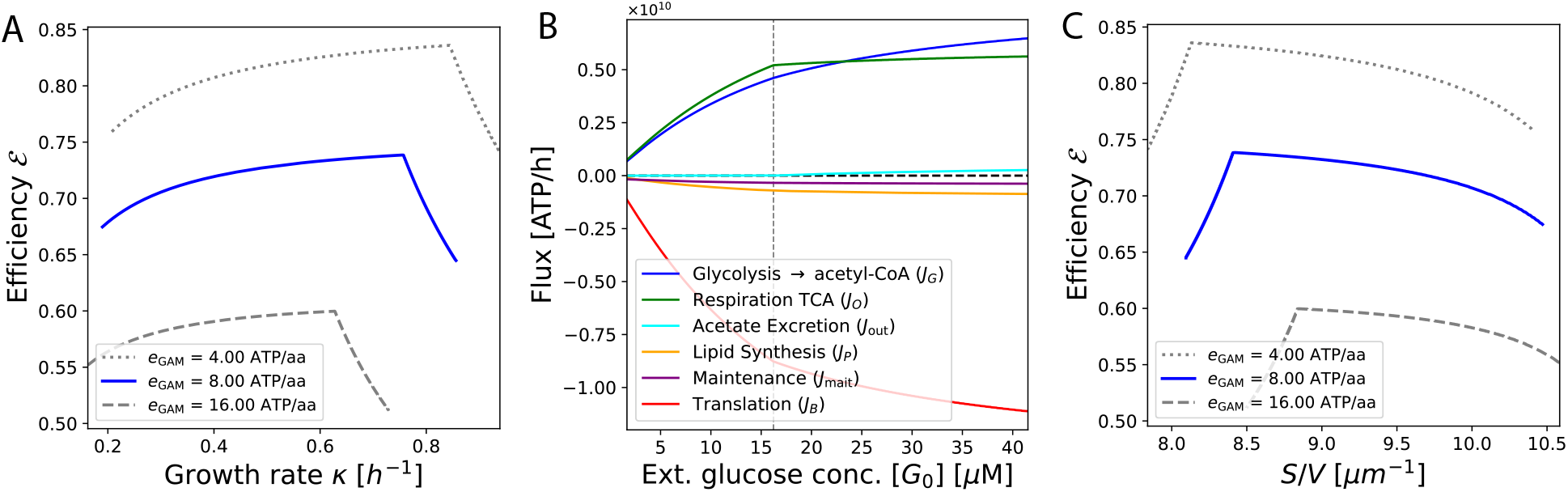
Optimal biosynthetic efficiency is achieved at the transition to overflow metabolism. (A) Biosynthetic energy efficiencies ℰ of our metabolic flux balance model, as defined in Eq. (5), as a function of growth rate and plotted parametrically for different values of the growth-associated maintenance *e*_GAM_ by varying the external nutrient concentration [*G*_0_]. In each case, efficiency increases up to the corresponding overflow transition point before dropping sharply: when the respiratory proteome is saturated, the cell trades biosynthetic efficiency to access higher growth rates. (B) With *e*_GAM_ = 8 ATP/aa, a breakdown of the ATP flux balance Eq. (4), illustrating the relative scale of each ATP production (positive flux) and consumption (negative flux) term in the equation. We vary the external glucose concentration [*G*_0_] on the x-axis, denoting the overflow transition with a vertical line. (C) Biosynthetic energy efficiency as a function of surface-to-volume ratio, plotted parametrically by varying the external nutrient concentration [*G*_0_].

As the cell transitions to overflow metabolism (shown for different values of *e*_GAM_ in Fig. 3A), the biosynthetic efficiency peaks and begins a sharp decline. This drop is a direct consequence of the onset of the waste cost (*J*_out_). The extra acetyl-CoA produced by glycolytic processes to meet ATP needs is excreted as acetate during fermentation, foregoing respiratory ATP. Thus, biosynthetic energy efficiency decreases asymptotically towards the theoretical minimum value; for a large enough cell in a rich enough environment, the lower limit on ℰ in overflow metabolism is the case where all energy is produced through fermentation (via acetate excretion).

We also observe that a larger growth-associated maintenance energy *e*_GAM_ leads to a decrease in the biosynthetic efficiency, since this results in less energy being allocated directly toward protein biomass growth. Interestingly, after the overflow transition, the subsequent decreases in ℰ seem to be of the same relative magnitude as the increases in ℰ observed in the non-overflow regime over the range of growth rates.

A breakdown of the ATP budget (Fig. 3B) reveals that protein translation is the dominant energy cost, an order of magnitude greater than maintenance or phospholipid synthesis. The budget also illustrates the shift in energy sourcing: in normal metabolism, glycolytic flux per hour (through the entire pathway in Fig. 1 resulting in acetyl-CoA) and respiration’s TCA cycle contribute roughly equally, but in overflow, the surface-limited respiratory flux flattens while glycolytic flux continues to increase to meet demand. Interestingly, while phospholipid synthesis has a high cost in terms of acetyl-CoA, the ATP costs are not a significant metabolic burden.

The relationship between biosynthetic energy efficiency and the surface-to-volume (*S/V*) ratio is also revealing (Fig. 3C). Counter-intuitively, the model predicts that the cells with the highest *S/V* ratio are the least energy efficient. This is a direct consequence of the empirical growth laws: the highest *S/V* cells are the smallest and slowest-growing, and their biosynthetic energy efficiency is thus dominated by the high relative cost of maintenance. Finally, it is important to note that our calculated energy efficiencies simplify the cell’s full ATP budget (e.g., ignoring DNA/RNA synthesis, protein degradation costs, etc.); including these additional metabolic loads would result in a more realistic energy efficiency. In addition, our model assumes that all the amino acids required for protein synthesis are synthesized solely from the carbon in the imported glucose, so the energy allocated for biosynthesis is notably higher than it would be in experiments where amino acids can be acquired from rich nutrient media.

### 3.3. Cell morphology shapes biosynthetic efficiency

The link between cell geometry and energy efficiency raises a key question: are the morphologies observed in nature optimal, or could cells achieve higher biosynthetic efficiency by adopting different forms? To answer this, we used our model to test the metabolic consequences of deviating from the empirically observed morphological scaling laws (Fig. 4).

**Figure 4.**
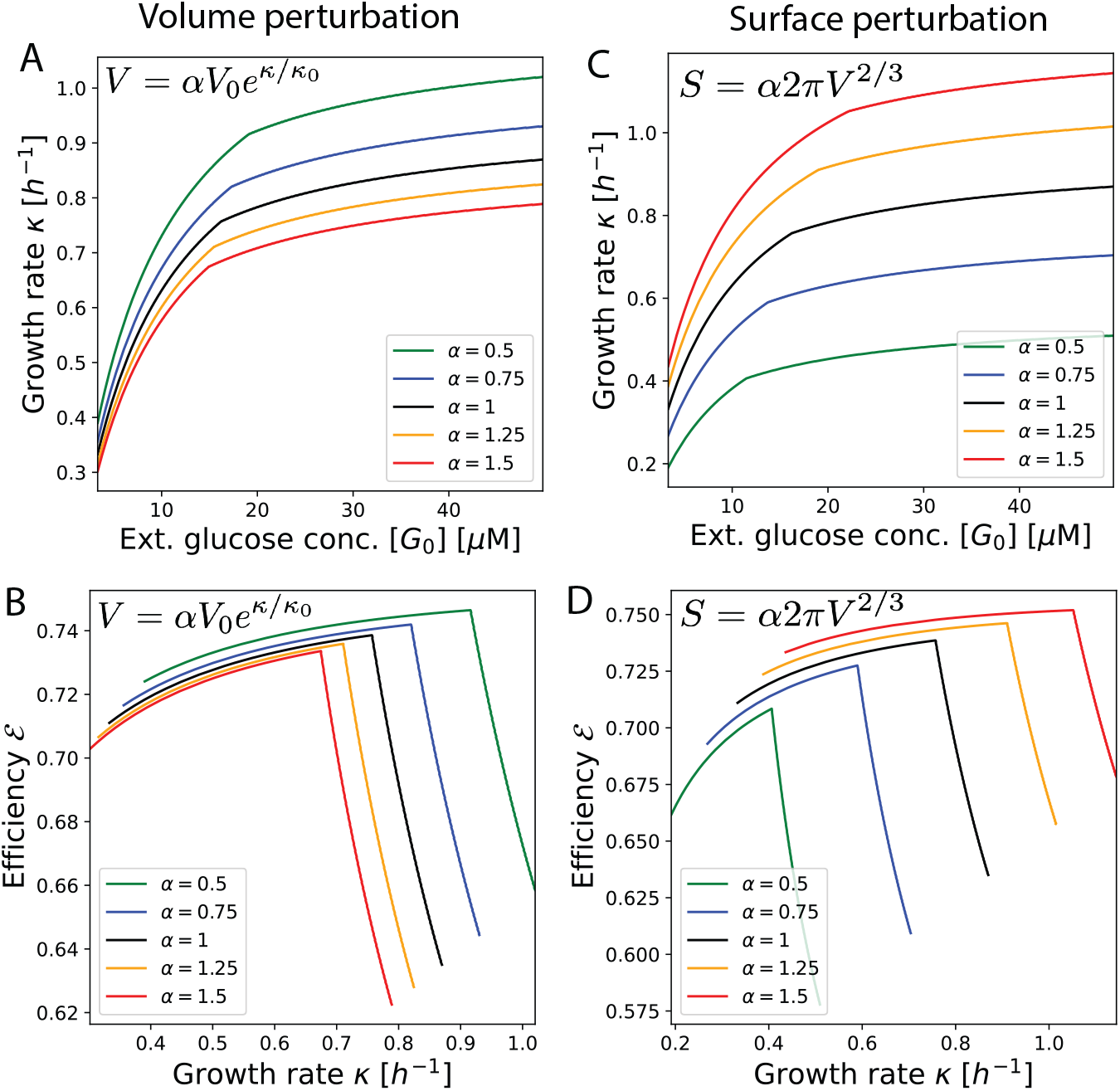
Cell morphology regulates biosynthetic efficiency. To test if observed cell morphologies are optimal, we simulated perturbations to cell volume and surface area. Each colored line represents a different morphological scaling law, controlled by the parameter *α*. (A, B) **Volume perturbations**. Here, we scaled the cell’s volume relative to the empirical growth law 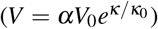 by a factor *α*. For *E. coli V*_0_ = 0.17 *µ*m^3^, *α* = 1 and *κ*_0_ = 0.86 h^−1^ [15]. (A) Increasing cell volume (*α >* 1) consistently reduces the growth rate *κ* at a given nutrient concentration. (B) Larger cells also exhibit lower biosynthetic efficiency ℰ at any given growth rate. (C, D) **Surface area perturbations**. Here, we scaled the cell’s surface area relative to its volume (*S* ∝ *αV* ^2*/*3^) by a factor *α*. (C) Increasing the surface area (*α >* 1) for a given volume increases the growth rate. (D) Cells with more surface area are also more energy efficient at any growth rate. Collectively, the results show that morphologies with a lower surface-to-volume ratio are metabolically disadvantageous.

First, we simulated cells that were constitutively larger or smaller than normal for a given growth rate (Fig. 4A-B). This is done by changing the parameter *V*_0_ in the relationship between cell volume *V* and nutrient-specific growth rate 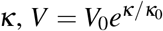. While bacterial cells tend to be larger at faster growth rates, our simulations show that artificially increasing cell size (by increasing *V*_0_) for any given nutrient condition is metabolically costly. Larger cells exhibit both a lower growth rate (Fig. 4A) and a lower biosynthetic efficiency ℰ (Fig. 4B). These larger cells also enter overflow metabolism at lower growth rates (Fig. 4B). This is a direct consequence of their reduced *S/V* ratio, which limits nutrient import and the capacity for efficient respiration, forcing them into overflow metabolism at lower growth rates. Conversely, constitutively smaller cells consistently achieve higher biosynthetic efficiency.

We found similar results when perturbing the surface area *S* for a given volume *V* (Fig. 4C-D). This is done by altering the cell shape factor *α* defined as *S* = 2*παV* ^2*/*3^, where *α* = 1 is the empirically observed value for *E. coli* [17]. Cells endowed with a higher surface area for a given volume (*α >* 1) grew faster and more efficiently. Taken together, these perturbations reveal a strong metabolic pressure to maximize the *S/V* ratio; any deviation toward a lower *S/V* ratio is metabolically disadvantageous.

This principle can be captured in a single, powerful analytical expression derived from our model. By solving the flux balance equations at the point of maximum efficiency (the onset of overflow) and using the parameters in Table I, we can find a direct relationship between the maximally efficient growth rate (*κ*_crit_) and the cell’s geometry (represented by the envelope fraction *ϕ*_*E*_):

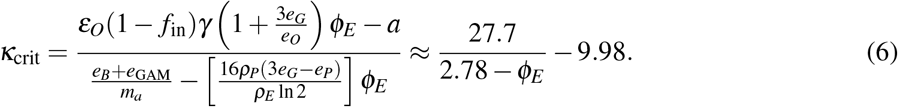

The above equation states that for any given cell geometry, there is a hard ceiling on the rate of efficient growth. *κ*_crit_ is a monotonically increasing function of *ϕ*_*E*_ (which is proportional to the surface area as 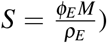, supporting the conclusion that when volume is held constant, an increase in the surface-to-volume ratio directly enables the cell to achieve a greater *maximally efficient growth rate* before overflow metabolism becomes necessary. When *κ*_crit_ = 0.76 h^−1^ [24], this equation predicts a 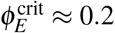, as can be observed in Fig. 2A.

### 3.4. Intracellular diffusion and crowding limit metabolism

Beyond the geometric constraints at the cell surface, a second physical challenge for metabolism is the time required for molecules to diffuse and react within the cytoplasmic volume. To capture this, we incorporated the *Internal Diffusion Constraint* hypothesis [35] by attenuating the rates of all internal metabolic fluxes (*J*_*G*_, *J*_*O*_, *J*_*B*_, *J*_*P*_) according to the characteristic intracellular diffusion time *T*. For a molecule diffusing across a cell of characteristic length *L* with diffusion constant *D*, this time scales as *T* ∝ *L*^2^*/D*.

Molecular crowding reduces the diffusion constant below its dilute-solution value as the cytoplasmic protein density rises. Following the scaled-particle treatment of Muramatsu and Minton [36], who showed that tracer diffusion in concentrated protein solutions decays exponentially with crowder concentration, we write

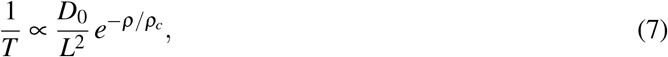

where *D*_0_ is the dilute-limit diffusion constant, *ρ* is the cytoplasmic protein density, and *ρ*_*c*_ is the characteristic cytoplasmic crowding density scale. This form captures two competing physical effects: the travel time grows with the square of the cell size, but is reduced in cells of lower cytoplasmic density through faster diffusion. We scale each internal metabolic flux by a factor *β/T* , normalized so that a reference cell with *ρ* = *ρ*_*c*_ experiences no net penalty relative to itself, and the fluxes are amplified or diminished only when *ρ* deviates from *ρ*_*c*_ (see Methods 5.2 for implementation). We exclude nutrient import (*J*_in_) and non-growth maintenance (*J*_mait_), which are not limited by intracellular diffusion.

As expected, including this diffusion limit reinforces the conclusions from the previous sections. Larger cells, with their lower *S/V* ratios, are now penalized twice: once for their limited surface area for efficient metabolism, and again for the increased diffusion time associated with a larger cell length. This combined effect further reduces their growth rate and efficiency.

This framework allows us to dissect the distinct metabolic roles of cell size and cytoplasmic density. In Figure 5, we perturbed the cytoplasmic density while keeping cell volume and surface area constant. With crowding switched off (*D* = *D*_0_, solid lines), less dense cells still grow faster and more efficiently (Fig. 5A–B). This may seem counterintuitive, since the surface-to-volume ratio is unchanged. However, a lower density means a lower total protein mass *M* for the same volume, which in our proteome-based model requires a larger envelope fraction (*ϕ*_*E*_) to enclose. This raises the surface-to-mass ratio, which is metabolically favorable. When crowding is included (Eq. (7)), the trend is magnified. Lowering the density now confers a double benefit, improving the surface-to-mass ratio while simultaneously raising the diffusion constant and accelerating internal reactions.

**Figure 5.**
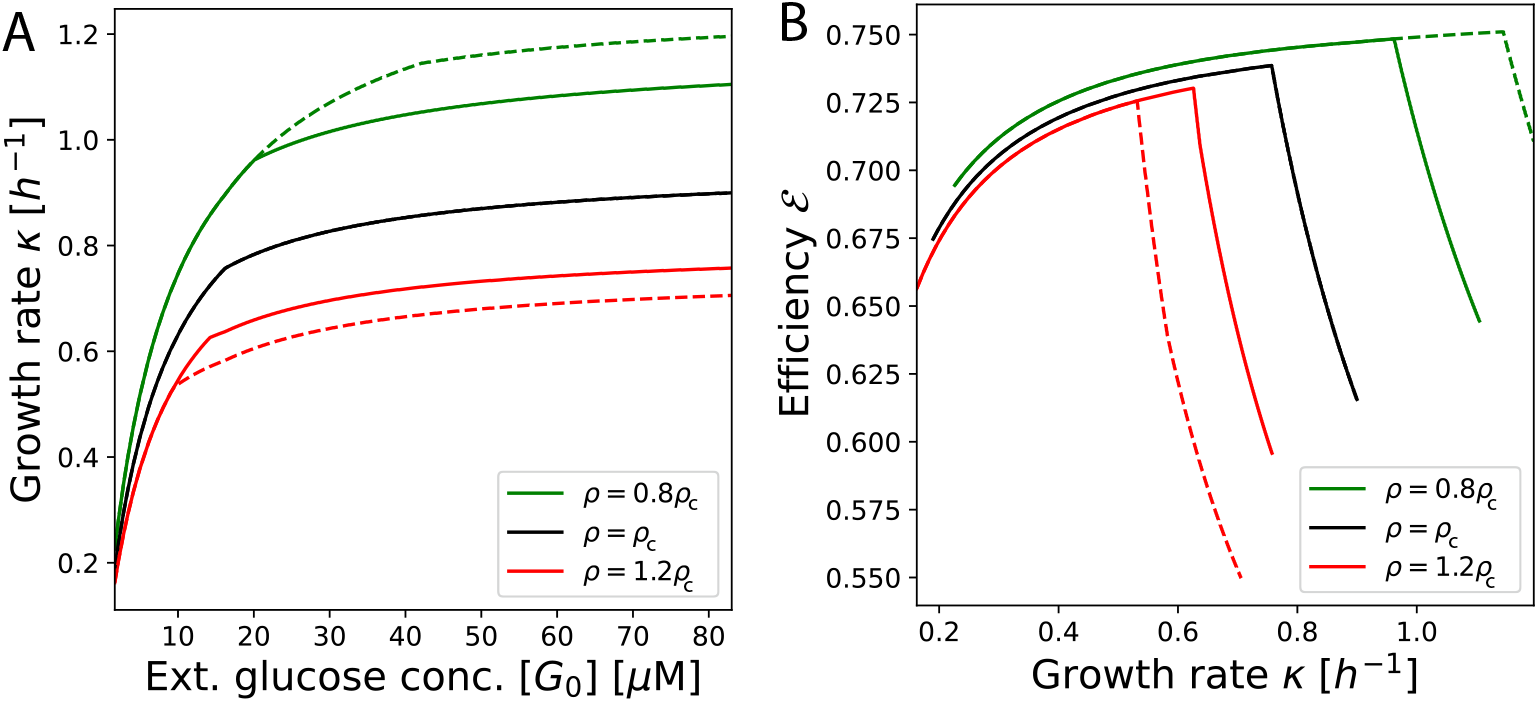
Intracellular diffusion and cytoplasmic density constrain cell growth and biosynthetic efficiency. The metabolic consequences of modeling intracellular diffusion and crowding. (A) Growth rate *κ* as a function of external glucose concentration, and (B) biosynthetic efficiency ℰ as a function of growth rate. Each color corresponds to a different cytoplasmic density (*ρ*) relative to the reference value (*ρ*_*c*_), where a change in density implies a change in protein mass for the same cell volume. Solid lines show the effect of diffusion limited by cell size only, while dashed lines include the additional effect due to internal density-dependent diffusion constraints. We scale the reaction rates by a factor *β/T* , where *T* is the characteristic diffusion time and *β* is a multiplicative constant that sets the normalization of the reaction rates to a reference value without diffusion limitations. The parameter *β* is fixed (see Eq. 17) for each value of [*G*_0_] such that *β/T* = 1 for *ρ* = *ρ*_*c*_. The results show that decreasing density improves both growth and efficiency, an effect that is magnified when crowding is considered.

These results reveal a trade-off governing reaction speed. A cell can be small and crowded (short diffusion path, but low diffusion constant *D*) or large and sparse (long path, but high *D*), with the overall reaction timescale *T* ∝ *L*^2^*/D* set by which factor dominates. Our model predicts that the optimal resolution of this trade-off is a cell that is both small and of low density. We note, however, that the model omits the opposing pressures that set the lower bounds on cell size and density, such as the physical volume required for the genome, stochastic effects, and the finite sizes of individual proteins [37]; it also coarse-grains the size dependence of crowding, applying a single crowding scale to all diffusing species, whereas in reality larger molecules are slowed more strongly [36].

### 3.5. Proteome budget crisis defines the upper limit of cell size

Having established the physical constraints that shape bacterial metabolism, we asked what sets the maximum sustainable cell size. We removed the constraint on surface area and volume imposed by the empirical growth laws and instead treated cell geometry as a free parameter, maximizing the volume *V* for a given target growth rate *κ* across a range of nutrient conditions. For simplicity we consider a spherical cell, parametrized by its radius *R* alone (see Methods 5.2 for implementation). Optimizing over the two independent shape parameters of a rod (radius and length) is substantially more involved and does not change the qualitative conclusion. What matters for the proteome budget crisis is that the intracellular diffusion length grows with cell volume, which holds for any cell that enlarges while preserving its shape.

The result is a striking inverse relationship: the maximum sustainable volume shrinks dramatically as the target growth rate rises (Fig. 6A), both with (dashed lines) and without (solid lines) diffusion limitation. A cell with negligible growth rate (*κ* = 0.0001 h^−1^) that need only cover maintenance can in principle reach volumes of ∼ 10^4^ *µ*m^3^ by relying on high nutrient import, whereas a cell growing at a modest *κ* = 0.5 h^−1^ is limited to ∼ 1 *µ*m^3^, which is a volume comparable to a typical bacterium.

**Figure 6.**
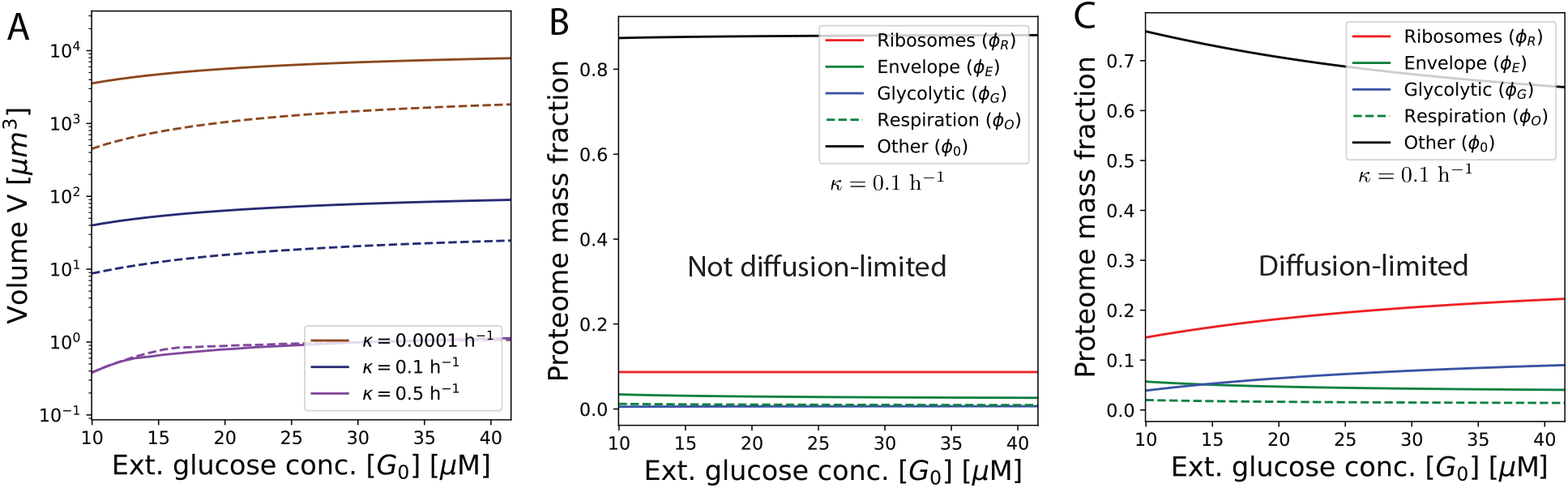
Theoretical maximum sustainable cell size is defined by a proteome budget crisis, with and without intracellular diffusion limitations. (A) Maximum sustainable volume *V* as a function of nutrient concentration when the empirical growth laws are relaxed. Cell protein mass *M* is used as a free parameter to maximize *V* = *M/ρ* for each value of [*G*_0_] (cytoplasmic protein density remains constant) at a chosen target growth rate. Solid lines show the limit set by nutrient import alone (the surface-to-volume bottleneck). Dashed lines show the much stricter limit imposed when intracellular diffusion is included. The results show an inverse relationship between growth rate and maximum cell size. (B, C) Proteome mass fractions as a function of external glucose concentration for a growing cell (*κ* = 0.1 h^−1^) without (B) and with (C) diffusion limits. Without diffusion, the proteome mass fractions remains largely unchanged. When diffusion is included, the “cost of size” becomes apparent: the ribosomal (*ϕ*_*R*_) and glycolytic (*ϕ*_*G*_) mass fractions must increase with [*G*_0_] to compensate for inefficiency, setting a hard limit on cell size.

This size limit arises from two compounding costs. First, a simple surface-to-volume ratio decrease in larger cells creates a nutrient import bottleneck. However, a more fundamental limit is imposed by an inescapable conflict in the proteome budget between the “cost of speed” and the “cost of size,” which becomes critical when intracellular diffusion is considered. The cost of speed is the high baseline investment in ribosomes (*ϕ*_*R*_) required for a fast growth rate. The cost of size is the additional investment in ribosomes and glycolytic enzymes (*ϕ*_*R*_, *ϕ*_*G*_) needed to overcome the inefficiency caused by longer intracellular diffusion paths in a larger cell. This effect is clearly visible in the proteome allocation (Fig. 6B, C). Without diffusion-limited reactions, most of the proteome does not need to be allocated to growth or metabolism to meet translation and ATP requirements even at large cell sizes (Fig. 6B). However, for the same growth rate with diffusion-limited reactions, the ribosome and glycolytic mass fractions must steadily increase with [*G*_0_], and hence the maximum sustainable volume (Fig. 6C). This is due to the increase in the diffusion length-scale; as translation and ATP synthesis become less efficient, they require a larger proteome investment.

A fast-growing cell, having already committed much of its proteome to the cost of speed, exhausts its remaining budget and cannot afford the cost of size, creating a proteome budget crisis that renders large volumes non-viable. The mechanism depends only on the intracellular diffusion length increasing with cell volume, and so is not specific to the cell geometry assumed here. It is this diffusion-driven proteome budget crisis, rather than nutrient import alone, that sets the upper bound on the size of a rapidly growing bacterium.

## 4. DISCUSSION

In this work we developed a mechanistic model to ask how a bacterium’s physical properties—its size, shape, and intracellular diffusion—constrain its metabolic strategy and growth. By integrating proteome allocation with the geometric limits of the cell surface and the biophysics of intracellular diffusion, our framework provides a first-principles account of several physiological observations. We demonstrated that overflow metabolism emerges as an optimal trade-off that bypasses the surface-area bottleneck on respiration to sustain rapid growth. Biosynthetic energy efficiency rises as respiration approaches its surface-limited capacity and peaks precisely at the onset of overflow; beyond this point, active overflow trades efficiency for growth rate, excreting carbon as acetate. The efficiency optimum therefore marks the boundary of the respiration-limited regime, while overflow itself is the growth-maximizing strategy the cell adopts once that boundary is reached. Together, these results recast a seemingly wasteful metabolic behavior as the rational consequence of a geometric limit on respiratory capacity.

The empirically observed growth laws relating cell size and shape to growth rate are not arbitrary in our model: they place cells close to the efficiency optimum set by the geometric advantage of a high surface-to-mass ratio, balanced against the costs of miniaturization. Whether this proximity reflects direct selection on efficiency or is a byproduct of other constraints is beyond the scope of the model, which contains no evolutionary dynamics; we show only that the observed morphologies sit near an efficiency optimum, not that efficiency is what selected them.

One of the key predictions of the model is an inverse relationship between growth rate and the maximum sustainable cell size. This ceiling is set not by nutrient import alone but by a conflict within the finite proteome budget. The cost of speed is the high ribosomal investment that fast growth requires; the cost of size is the additional investment needed to offset slower intracellular diffusion in a larger cell. For a fast-growing cell these become mutually exclusive, and the resulting proteome budget crisis makes large volumes non-viable. The mechanism depends only on the intracellular diffusion length increasing with cell volume, and so is not specific to the cell geometry we adopted for tractability; it offers a physical reason why rapidly dividing bacteria are constrained to remain small.

The model’s value lies in its simplicity, and that simplicity also bounds its claims. We do not account for all cellular ATP costs, such as DNA and RNA synthesis or protein turnover, so the absolute efficiency values are upper estimates and the true maximum is lower. We treat the proteome at a coarse-grained level and do not resolve regulation, which may explain why the predicted acetate flux rises nonlinearly with nutrient level where experiments show a more linear dependence on growth rate. Our treatment of intracellular diffusion also coarse-grains the size-dependence of crowding, applying a single crowding scale to all diffusing species, whereas larger molecules are in reality slowed more strongly. Even with these simplifications, the framework shows that a few physical and budgetary rules are sufficient to define the metabolic strategies and size range available to bacteria.

The principles identified here offer a foundation for exploring metabolic constraints in more complex cells. Eukaryotes generate ATP on internal mitochondrial membranes rather than the cell surface, relieving the surface-area bottleneck that governs bacterial respiration [38]. These membranes remain subject to crowding [39] but are no longer tied to the cell’s surface-to-volume ratio, and can be regulated separately from nutrient uptake. Eukaryotic cells nonetheless still exhibit overflow metabolism [40], the Warburg effect, and the relationship between ATP synthesis and eukaryotic cell size is not fully understood [41, 42]. Extending the present framework to capture organelle-based energy generation may help clarify the size scales and energetic trade-offs that govern both prokaryotic and eukaryotic life.

## 5. METHODS

### 5.1. Parameter determination

Most of our model parameters can be estimated from experimental data, and are collected in Table I. In this section, we calculate the parameters that are not directly measured or available from experiments.

#### Cell composition: protein and lipid densities

*E.coli* cells are approximately 55% protein by dry mass, so we can use a conversion factor of 0.55 to relate cellular dry mass to protein dry mass [21]. Assuming a uniform cell dry mass density of *ρ*_cell_ = 0.30 g/ml = 1.8 × 10^5^ MDa*/µ*m^3^ for *E. coli* [47], we can calculate the characteristic cytoplasmic protein dry mass density, *ρ*_*M*_, as:

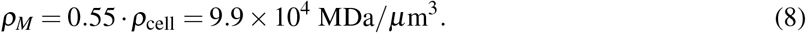

Given that the cell envelope has a thickness of *d* = 28.7 nm [46], we can determine the protein density per unit area of the cell envelope in a simplified way as *ρ*_*E*_ = *ρ*_*M*_*d* = 2.8 × 10^3^ MDa*/µ*m^2^. However, assuming that the protein mass density per area of the cell envelope is the same as that of the cytoplasm may not be warranted. Therefore, we convert experimental values [45] for the protein mass densities per area in the inner membrane and use *ρ*_*E*_ = 2.4 × 10^3^ MDa/*µ*m^2^ for our simulations, a relatively close value to our simple estimate.

#### ATP synthesis and overflow threshold

The cell’s respiratory energy production is largely driven by membrane-bound ATP synthase and electron transport chain complexes. A single ATP synthase has a molecular mass of *m*_ATPase_ = 545 kDa [48] and is estimated to occupy a surface area of *A*_ATPase_ = 35 nm^2^ on the inner membrane. When coupled with the electron transport chain (ETC) complex, this occupies *A*_ETC-ATPase_ = 84 nm^2^ in area [26]. We estimated the mass of a representative ETC complex in *E. coli* using the multi-subunit complexes named in [26]. For each complex, we used Gemini LLM [49] to map its constituent sequences to their respective UniProt [50] identifiers and to obtain the corresponding sequence-based masses. These aggregated masses were then multiplied by the fractional stoichiometries for each subunit complex defined in Fig. B1/Table S1B of Szenk et al. [26] to yield the final representative value. This resulted in the form 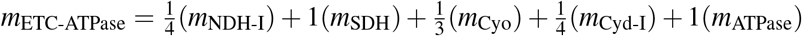, quantified as 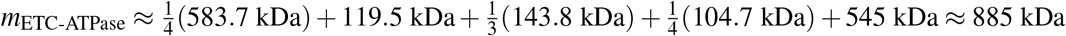. (An independent estimate where the masses were aggregated using Claude LLM [51] yielded ≈ 873 kDa; we use the former value and accept the marginal difference within tolerable limits for our coarse-grained estimations.) Fig. 2A of [26] shows that when these ETC complexes occupy about 9% of the total inner membrane surface area, the respiratory chain is saturated, and overflow metabolism sets in. This physical constraint allows us to define the maximum fraction of envelope protein mass dedicated to membrane-bound respiratory complexes, *ϕ*_*O*,max_, in relation to the total envelope protein fraction, *ϕ*_*E*_:

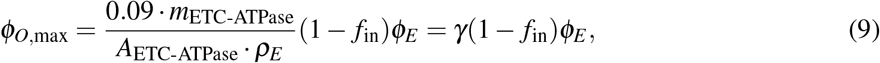

Here, the coefficient *γ* is calculated to be 0.39.

#### Proteome efficiencies

With each ATP synthase complex generating approximately 270 ATP molecules per second [52], the specific ATP production rate from respiration, *ε*_*O*_, per unit of protein mass is:

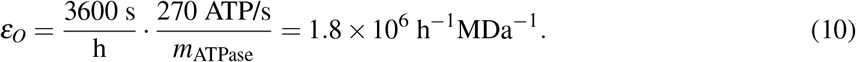

Since our model includes the glycolytic pathway (import of carbon to its conversion to pyruvate, and pyruvate to acetyl-CoA) as common to respiration and fermentation, and allows the final fermentative step to be encoded in *J*_out_ entirely, here we rescale the relation for *ε*_*F*_ = 1.92*ε*_*O*_ in [24] to obtain a proteome efficiency for *ε*_*G*_. The full fermentative pathway produces 12 ATP from one molecule of glucose. We only consider the glycolytic pathway (including NAD+ and NADH steps in Extended Data Figure 2 of [24] for oxidative fermentation) and exclude the 2 ATP from acetate excretion. Using a stoichiometric approach, we represent the efficiencies as

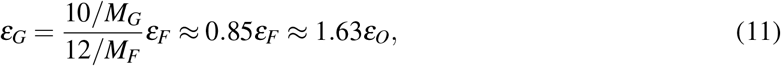

where *M*_*G*_ is the total mass of all the proteins involved in the glycolytic chain (glycolysis proteins and pyruvate dehydrogenase complexes to obtain acetyl-CoA), and *M*_*F*_ is the total mass of all the proteins involved in the entire fermentative pathway (encompassing *M*_*G*_ as well as the proteins that convert acetyl-CoA to acetate). These masses were computed from quantitative *E. coli* proteomics data (Table S6 of Supplementary Data in Schmidt et al. [9]) using Claude LLM [51] to extract all the proteins associated with the corresponding functional sectors, and multiplying the respective protein copy numbers per cell with the masses per protein. The cumulative protein masses involved in the glycolytic and pyruvate dehydrogenase pathway totaled *M*_*G*_ ≈ 9.10 × 10^6^ kDa, and the cumulative protein masses involved in the full fermentative pathway totaled *M*_*F*_ ≈ *M*_*G*_ + *M*_acetate_ ≈ 9.32 × 10^6^ kDa. It is worth noting that the small proteins involved in converting acetyl-CoA to acetate—*pta* and *ackA*—comprise only 2.4% of *M*_*F*_ (i.e. *M*_acetate_ ≈ 0.22 × 10^6^ kDa) despite allowing for 2 out of the 12 ATP per glucose to arise from acetate excretion, indicating that although overflow metabolism is carbon-inefficient, it is highly proteome-efficient.

#### Lipid synthesis and stoichiometry

Lipid synthesis is another crucial anabolic process, starting from the precursor acetyl-CoA [53]. The creation of a single C16 fatty acid (palmitate) requires 8 acetyl-CoA molecules and 21 ATP equivalents. These fatty acids are then assembled into phospholipids, which are the primary components of cell membranes [44]. Therefore, taking palmitate as a representative acyl chain, the synthesis cost per acetyl-CoA incorporated is *e*_*P*_ = 21*/*8 ATP/acetyl-CoA.

To estimate the area density of phospholipids, we consider their physical properties. Each phospholipid occupies an area of *A*_*P*_ = 0.65 nm^2^ and constitutes about 50% of the membrane’s mass. With a mass of 0.691 kDa and a volume of 1.25 nm^3^ [54], a phospholipid’s density is ∼0.92 g/cm^3^, compared to the average protein density of 1.35 g/cm^3^ [55]. Using these values, the phospholipid area density, *ρ*_*P*_, is estimated as:

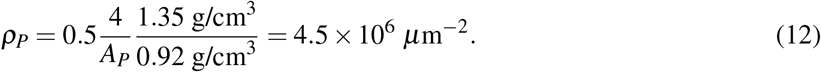

The factor of 4 accounts for the two leaflets (bilayers) of both the inner and outer membranes. Since each phospholipid incorporates 16 acetyl-CoA (owing to its two fatty acid chains), the acetyl-CoA flux directed toward lipid synthesis, *J*_*P*_, must remain balanced with protein synthesis over the cell cycle duration, *τ*:

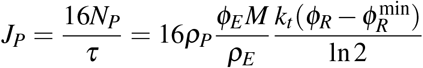

Here, *N*_*P*_ is the initial number of phospholipids in the cell.

#### Physical limits on cell morphology

While resource allocation imposes an upper limit on the fraction of protein dedicated to the cell envelope, *ϕ*_*E*_ , a physical lower limit also exists. This minimum is dictated by the geometry of a sphere, which represents the smallest possible surface area for a given cellular volume. The surface area, *S*, of a spherical cell can be expressed in terms of its total protein mass *M* and protein mass density *ρ*:

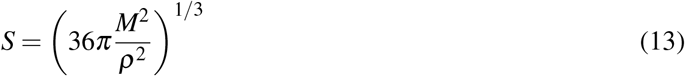

Using the relation above in conjunction with *S* = *ϕ*_*E*_*M/ρ*_*E*_ and *ρ* = *ρ*_*M*_, we can derive the minimum possible envelope protein allocation, *ϕ*_*E*,min_, for a viable spherical cell:

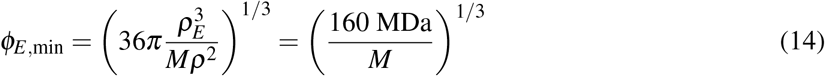

This equation shows that the minimum required envelope fraction is inversely related to the cell’s total protein mass.

#### Conversion of GAM to units of e_GAM_

While the exact values of growth-associated maintenance (GAM) energies can be calibrated based on specific constraints, here we compute a baseline approximation from reported data for GAM for *E. coli* under conditions of aerobic growth on glucose. Feist et al. [32] reports GAM = 59.81 mmol ATP gDW^−1^, representing ATP hydrolyzed per gram of new biomass formed. The amino-acid content of biomass follows from a protein fraction of *f*_*p*_ = 0.55 g gDW^−1^ and a mean residue mass of *m*_*a*_ = 110 g mol^−1^, giving *n*_*aa*_ = *f*_*p*_*/m*_*a*_ = 5.0 mmol gDW^−1^, the molar quantity of residues per gram dry weight. Dividing the GAM by this content yields 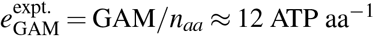. However, because the reported value of GAM in [32] already includes the 4 ATP aa^−1^ consumed to polymerize residues during protein translation (already counted in *e*_*B*_), we subtract this to avoid double-counting, leaving an effective

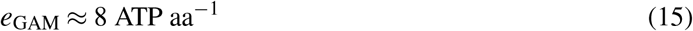

as the net growth maintenance overhead. For comparison, *e*_*B*_ = 23.5 ATP aa^−1^ captures the full biosynthesis and translation cost, including the opportunity cost of diverting precursors from energy production [33, 56].

#### Proteome allocation of glucose transporters

Consider an *E. coli* unit cell defined by volume *V* = 1 *µ*m^3^ and inner membrane surface area *S* = 6 *µ*m^2^ [57]. The total area occupied by PtsG glucose transporters, *A*_ptsg_, is determined by the copy number *n*, the cell volume *V* , and the cross-sectional area of the protein dimer *a*_ptsg_, as obtained from [58]:

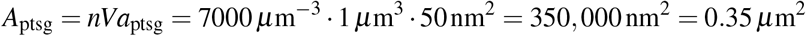

The total area available to the membrane proteome, *A*_prot_, is defined by the surface area *S* and the fractional protein occupancy *f*_prot_ of all membrane proteins relative to the total surface area (60% based on [45]):

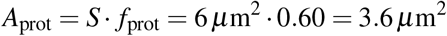

*ϕ*_in_ represents the mass fraction of PtsG glucose transporters relative to the total envelope proteome mass fraction *ϕ*_*E*_. Assuming the mass-to-area density of PtsG is consistent with the average of the membrane proteome (not unreasonable for transmembrane proteins), we have the ratio of these areas:

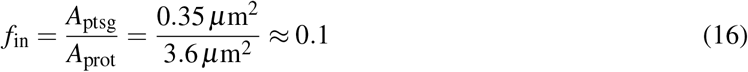

This establishes the final relationship for the PtsG mass fraction:

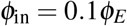

Thus, glucose transporters account for approximately 10% of the total mass of the membrane envelope proteome.

### 5.2. Modeling intracellular diffusion

Cell volume and mass are known to decouple in long-term evolution experiments; cells tend towards decreasing mass-to-volume ratio [35]. One such explanation for this phenomenon is the *Internal Diffusion Constraint* hypothesis: cellular metabolism is limited by cytoplasmic crowding and intracellular diffusion length scales. The larger and less dense cells seen in these experiments have more surface area with which to import more nutrients and faster intracellular diffusion because of a decrease in crowding. Previous work analyzing these experiments has considered that metabolic reaction rates scale with the reciprocal of the average travel time between reactants 1*/T* [35]. For a diffusing particle, distance traveled scales with the square root of time, i.e., *T* ∝ *L*^2^*/D* for a spherical cell, where *L* is cell length and *D* is the diffusion constant.

To apply intracellular diffusion constraints, we scale the fluxes *J*_*G*_, *J*_*O*_, *J*_*B*_, and *J*_*P*_. We exclude *J*_in_ and *J*_mait_ since these fluxes correspond to external diffusion and non-growth associated maintenance. We implement the scaling factor discussed in Eq. (7) by solving for a proportionality constant *β* ^′^ in

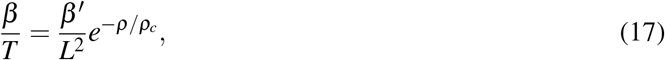

where *β* ^′^ absorbs the diffusion constant *D*_0_ seen in Eq. (7). Absent an independent measurement of *ρ*_*c*_ in *E. coli* systems, we assume *ρ*_*c*_ = *ρ*_*M*_, which places the reference cell at *e*^−1^ of its dilute-limit diffusion constant. Using Eq. (17), we scale the metabolic fluxes of our cell by multiplying them by *β/T* , where the diffusion-encoding *β* ^′^ is solved for each [*G*_0_] such that *β/T* = 1 when *ρ* = *ρ*_*c*_ = *ρ*_*M*_. This establishes the reference baseline flux for which there is no diffusive penalty. When the internal diffusion constraint is applied, changes in cytoplasmic protein density lead to non-linear changes in the diffusion time for metabolites. When *ρ > ρ*_*c*_, the scaling factor decreases, suppressing diffusion and growth rate. When *ρ* < *ρ*_*c*_, the scaling increases, enhancing both. As shown in Fig. 5, equal magnitudes of Δ*ρ* = *ρ* − *ρ*_*c*_ about *ρ*_*c*_ produce asymmetric responses: the growth rate shift is larger when Δ*ρ* < 0.

For rod-like cells (as in Fig. 5), we model *L* as the end-to-end length of the cell including the hemispherical caps. The cell surface and volume, *S* = 2*πR*(*L* − 2*R*) + 4*πR*^2^ = 2*πRL* and *V* = *πR*^2^*L* − 2*πR*^3^*/*3, give two equations for the unknowns *R* and *L*. Substituting *L* = *S/*(2*πR*) yields a cubic in *R*, which we solve numerically; we then compute *L* = *V/*(*πR*^2^) + 2*R/*3. We implement the intracellular diffusion length as *L*_diffusion_ = (*L* − 2*d*) to account for envelope thickness on both ends of the cell axis. For spherical cells (as in Fig. 6), the diffusion length is given by cell radius as *R*_diffusion_ = *R* − *d* = 3*V/S* − *d*, with a single envelope thickness subtracted for the radial path. Therefore, for both rod-like and spherical geometries, *R* is set by *S* and *V* , which follow from *ϕ*_*E*_ , *ρ*_*E*_ , *M*, and *ρ*.

### 5.3. Calibration with bacterial growth laws and flux balance equations

We calibrate our model with established empirical data on bacterial growth. We take the dependence of cell volume on growth rate, *V* (*κ*) = 0.17*e*^1.16*κ*^ *µ*m^3^ (with *κ* in h^−1^) [15], and the surface-area-to-volume scaling law in *E. coli, S* = 6.24*V* ^2*/*3^ *µ*m^2^ [17]. Combined with the cytoplasmic protein density *ρ*_*M*_ and the envelope protein density *ρ*_*E*_ , these give the total protein mass *M* = *ρ*_*M*_*V* and the envelope protein fraction *ϕ*_*E*_ = *ρ*_*E*_*S/M* for any growth rate *κ*. Because our model links *κ* to the ribosomal protein fraction *ϕ*_*R*_ (Eq. (1)), and *V* to *κ* through the growth law above, both *M* and *ϕ*_*E*_ are determined by *ϕ*_*R*_.

We calibrate *K*_*C*_ as a free parameter used to scale the nutrient influx capacity via porins in the term *J*_in_. We constrain this with the experimental observation that overflow metabolism in *E. coli* begins at *κ* = 0.76 h^−1^ (fitted over various *E. coli* strains and nutrients such as glucose [24]). Choosing *K*_*C*_ = 30 *µ*M and using our derived estimate for the growth-associated maintenance energy *e*_GAM_ = 8 ATP/aa in Eq. (15), we obtain an overflow transition around *κ* = 0.76 hr^−1^. A parameter sensitivity analysis for *K*_*C*_ is shown in Fig. S1, demonstrating that *K*_*C*_ has a negligible influence on the transition growth rate and maximal efficiency at the overflow transition, though it does affect the nutrient concentration at the transition point (when *K*_*C*_ approaches the glucose transporter’s *K*_ptsG_ = 10 *µ*M, a larger [*G*0] is required to achieve overflow).

At the overflow transition point, the respiratory system is maximally utilized 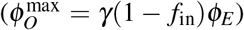, while acetate secretion is zero (*J*_out_ = 0), after which the respiratory proteome is capped at 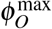 and *J*_out_ is solved for. The flux balance equations for carbon, acetyl-CoA, and ATP in Eqs. (2), (3), and (4) involve four quantities that can vary (*ϕ*_*R*_, *ϕ*_*G*_, *ϕ*_*O*_, *J*_out_); the overflow regime fixes one of these, leaving three equations for three unknowns. Below the overflow transition, respiration is unsaturated and *J*_out_ = 0, so we solve for *ϕ*_*R*_, *ϕ*_*G*_ and *ϕ*_*O*_. Above the transition, *ϕ*_*O*_ is held at 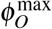 and the surplus acetyl-CoA is excreted as acetate, so we solve for *ϕ*_*R*_, *ϕ*_*G*_ and *J*_out_.

### 5.4. Critical condition for overflow metabolism

We aim to find a relationship between the cell’s envelope proteome fraction, *ϕ*_*E*_ , and the critical growth rate, *κ*_crit_, at which the cell transitions to overflow metabolism. At this transition point, two conditions are met: (1) Acetate excretion flux is zero: *J*_out_ = 0. (2) The respiratory proteome is saturated: 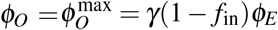, where *γ* = 0.39. We begin with the steady-state flux balance equations for acetyl-CoA and ATP, divided by the total cell protein mass *M* to yield specific fluxes (*j*_*X*_ = *J*_*X*_ */M*). The steady-state acetyl-CoA balance is given by:

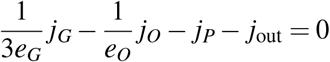

At the overflow transition point (*j*_out_ = 0), and this simplifies to:

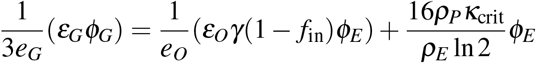

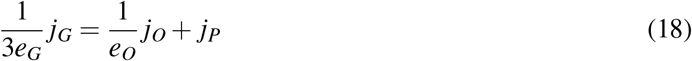

We substitute the definitions for the specific fluxes: *j*_*G*_ = *ε*_*G*_*ϕ*_*G*_, *j*_*O*_ = *ε*_*O*_*ϕ*_*O*_ = *ε*_*O*_*γ*(1 − *f*_in_)*ϕ*_*E*_ , and 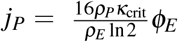.

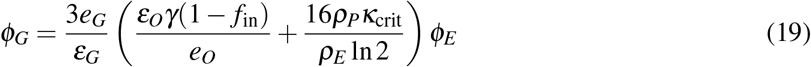

This equation links the necessary glycolytic proteome fraction *ϕ*_*G*_ to the envelope fraction *ϕ*_*E*_ at the critical growth rate *κ*_crit_.

The steady-state ATP balance is given by:

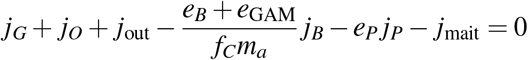

With *j*_out_ = 0, *j*_*B*_ = *f*_*C*_*κ*_crit_, and *j*_mait_ = *a*, this becomes:

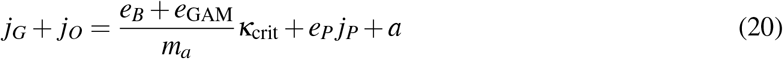

Substituting the flux definitions yields:

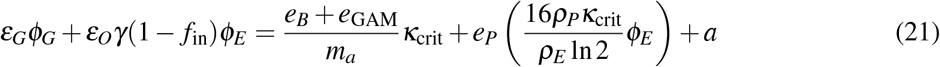

We now substitute the expression for *ϕ*_*G*_ from Eq. (19) into the ATP balance Eq. (21):

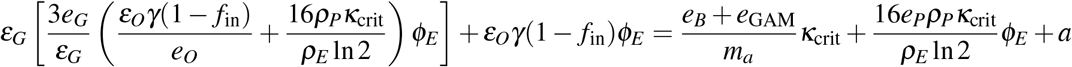

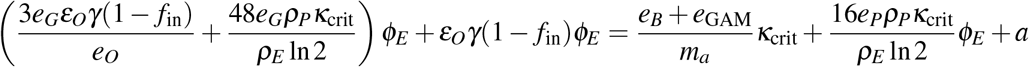

Next, we group all terms containing *ϕ*_*E*_ on the left-hand side:

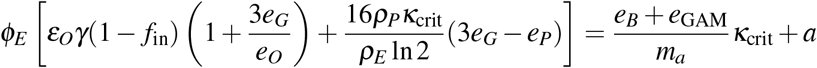

Finally, solving for *ϕ*_*E*_ yields the analytical condition for the onset of overflow metabolism:

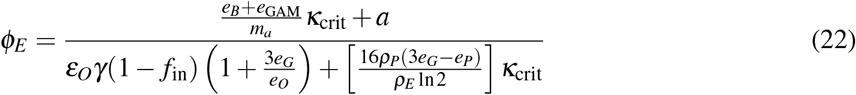

This equation provides a direct link between the cell’s envelope proteome fraction (*ϕ*_*E*_) and the maximum growth rate it can achieve (*κ*_crit_) at the time of the overflow transition.

## Acknowledgments

S.B. acknowledges support from the National Institutes of Health (NIH R35 GM143042), and the Shurl and Kay Curci Foundation.

## Author Contributions

AC and SB conceived and designed the research. AC, IB and SB developed the theoretical model. AC and IB performed simulations and data analysis. AC, IB and SB wrote the paper.

---

**Figure S1.**
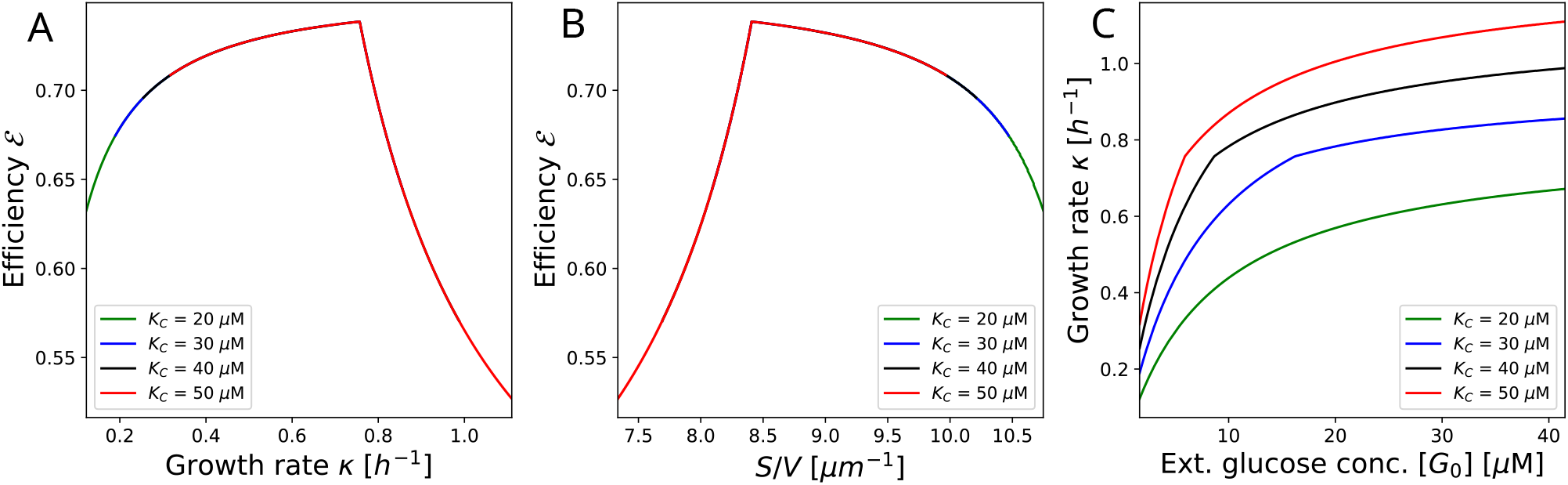
Effects of varying nutrient influx capacity via the porin-associated free parameter *K*_*C*_ on biosynthetic efficiency and growth rate. (A, B) Biosynthetic efficiency ℰ as a function of growth rate *κ* (A) and surface-to-volume ratio (B), plotted parametrically over a range of glucose concentrations. In each case, the efficiency increases as a function of growth rate and *S/V* up to the overflow transition point, after which the efficiency drops. The specific growth rate at which the overflow transition occurs is negligibly affected by an increase in *K*_*C*_, occurring for each tested *K*_*C*_ at ≈ *κ* = 0.76 hr^−1^. Since the porin influx term in *J*_in_ scales proportionally as *P*^′^ = *PK*_*C*_, this demonstrates that increasing the influx channel capacity does not significantly affect the transition growth rate and maximal efficiency at which the switch to overflow metabolism occurs. (C) Growth rate *κ* as a function of the external glucose concentration [*G*0]. For the lowest *K*_*C*_, the overflow transition does not occur over the plotted range of [*G*0]; however, for *K*_*C*_ = (30, 40, 50) *µ*M, the overflow occurs at the same growth rate *κ* ≈ 0.76 hr^−1^, but differing glucose concentrations. Increasing the nutrient influx capacity causes the cell to transition to overflow metabolism at a lower nutrient concentration, and achieve a greater growth rate for a given [*G*0].

## Notes

### Competing Interest Statement

The authors have declared no competing interest.

